# A Practical Roadmap For Sampling Floral Nectar From Communities of Many Plant Species

**DOI:** 10.64898/2025.12.19.695174

**Authors:** G.E. Kirschke, J.A. Bain, J.E. Ogilvie, P.J. CaraDonna

## Abstract

1. Floral nectar plays a critical role in shaping the ecology and evolution of plant-pollinator interactions. Effective and efficient methods that allow for broad-scale sampling of nectar volume and sugar concentration across a diversity of taxa are needed to improve our understanding of many dimensions of mutualistic plant-pollinator interactions—including their basic ecology and evolution, their responses to environmental change, and their conservation and restoration.
2. Despite the key importance of nectar for mediating plant-pollinator interactions, quantifying floral nectar in the field from many different plant species is challenging because there is often no one-size-fits-all sampling method that is effective across a diversity of floral structures and nectar traits. Different methods require different preparation, and sampling from many species involves a variety of logistical challenges.
3. Here we provide a methodological roadmap for sampling floral nectar in the field from many different plant species. We describe our nectar collection methods in detail, including necessary equipment, calculations, and approaches appropriate for different floral morphologies. We also provide a troubleshooting guide for common problems encountered while collecting nectar in the field. To demonstrate the utility and effectiveness of our methods for collecting nectar from many different species, we present results on nectar trait variation from 53 species in an ecosystem.
4. Our method illustrates that nectar traits vary considerably within and among plant species, indicating that large-scale nectar sampling projects are an important consideration for many basic and applied questions in pollination ecology and evolution. We hope that across many plant communities and ecosystems, our paper provides a practical roadmap for how to navigate the complexities of quantifying floral nectar traits.

## 1. Introduction

### 1.1 Background

At the center of mutualistic interactions among plants and pollinators is an exchange of floral food resources for assistance with sexual reproduction. Floral nectar shapes plant-pollinator interactions by attracting pollinators to facilitate plant reproduction and providing an important food source for many pollinators (Wilmer, 2011). Floral nectar is a complex and dynamic resource, with nectar traits—volume, sugar concentration, total sugars (among others)—varying widely across plant species (Nicolson, 2022; Figure 1). Despite the importance of nectar for mediating the mutualistic interactions between plants and pollinators, relatively few community-level studies have quantified nectar traits beyond single species or a small subset of species within a community (e.g., Baude et al., 2016; Fantinato et al., 2021).

**Figure 1.**
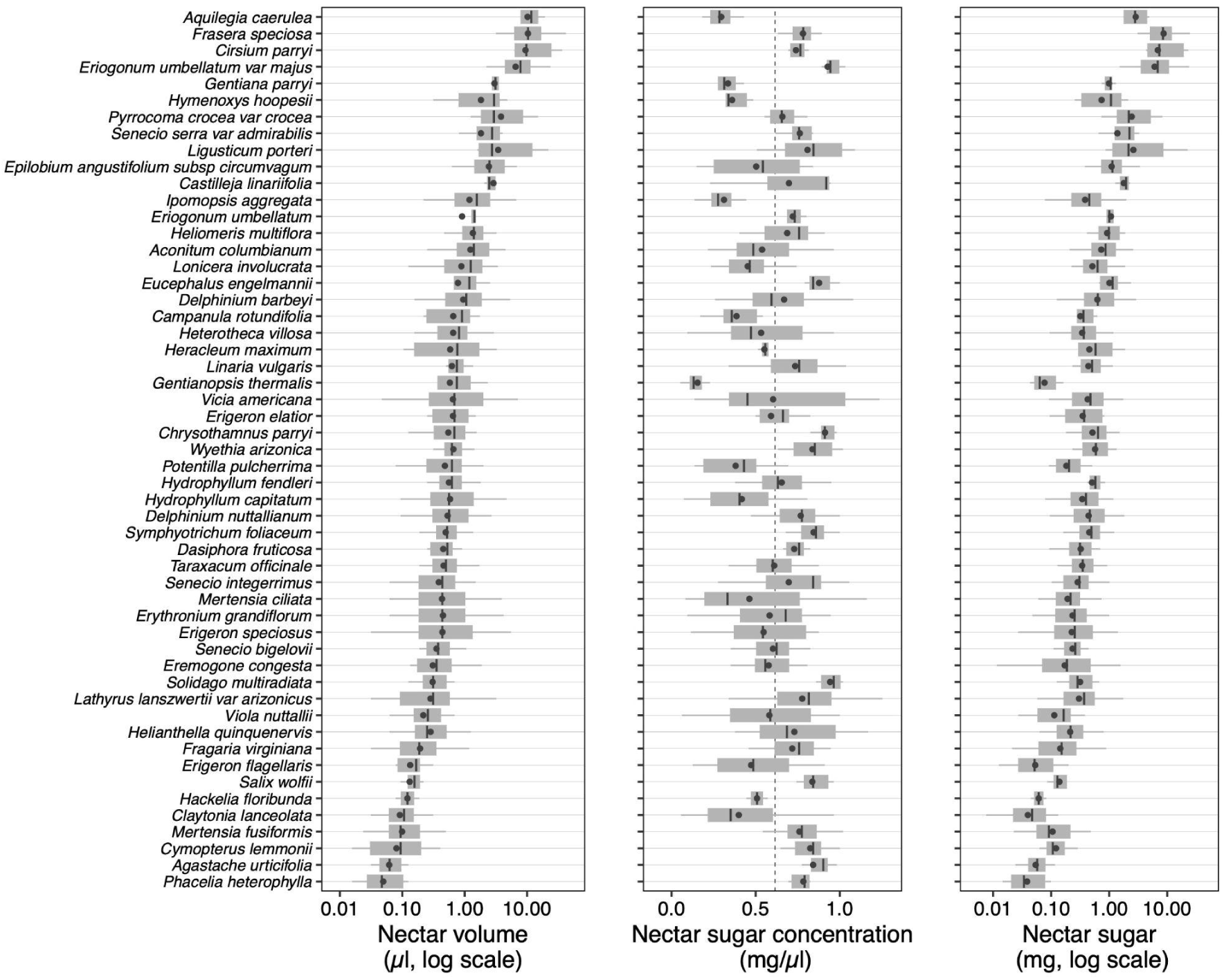
Nectar traits for the 53 plant species sampled in a montane plant community in Gothic, Colorado, USA. Black dots indicate mean nectar values; thick black lines represent the median; gray boxes represent the 25th and 75th percentiles; and the whiskers demonstrate the minimum and maximum values. The dashed line on the nectar concentration panel indicates 50% Brix, which is a cutoff point for which sugar refractometer to use when sampling nectar in the field (see Section 3.5). Note that nectar volume and nectar sugar are both plotted on the log scale to account for the wide range of values measured across species.

Effective and efficient methods that allow for broad-scale sampling of nectar traits across a diversity of taxa are needed to improve our understanding of many dimensions of mutualistic plant-pollinator interactions—including their basic ecology and evolution, their responses to environmental change, and their conservation and restoration. For example, nectar sampling across many species is critical for understanding spatiotemporal variation in carbohydrate availability for pollinators, or plant energetic investment in competition for pollination. General methods for floral nectar collection are available (*reviewed in* Kearns & Inouye, 1991), but quantifying floral nectar in the field from many different plant species is challenging for a few, interrelated reasons: first, there is often no one-size-fits-all sampling method that is effective across a diversity of floral structures and nectar traits in a given community; second, different methods require different preparation; and third, applying different techniques to a large number of species over an entire growing season poses logistical challenges.

In this paper we provide a practical roadmap of floral nectar collection techniques and how to apply them to a new field system. The techniques presented are effective for quantifying nectar volume (μl/flower), sugar concentration (mg/μl), and total sugars (mg/flower) for many different plant species (e.g., Figure 2), and we demonstrate the utility of these methods across 53 plant species over 8 years. Our nectar collection roadmap comes in three parts: (1) a brief review of nectar collection methods and results from their application in a subalpine system, (2) a discussion of factors affecting planning and workflow, and (3) a troubleshooting guide for common problems encountered in the field. The troubleshooting guide can assist in applying methods to new contexts. A detailed overview of planning considerations and nectar collection methods can be found in the appendix.

**Figure 2.**
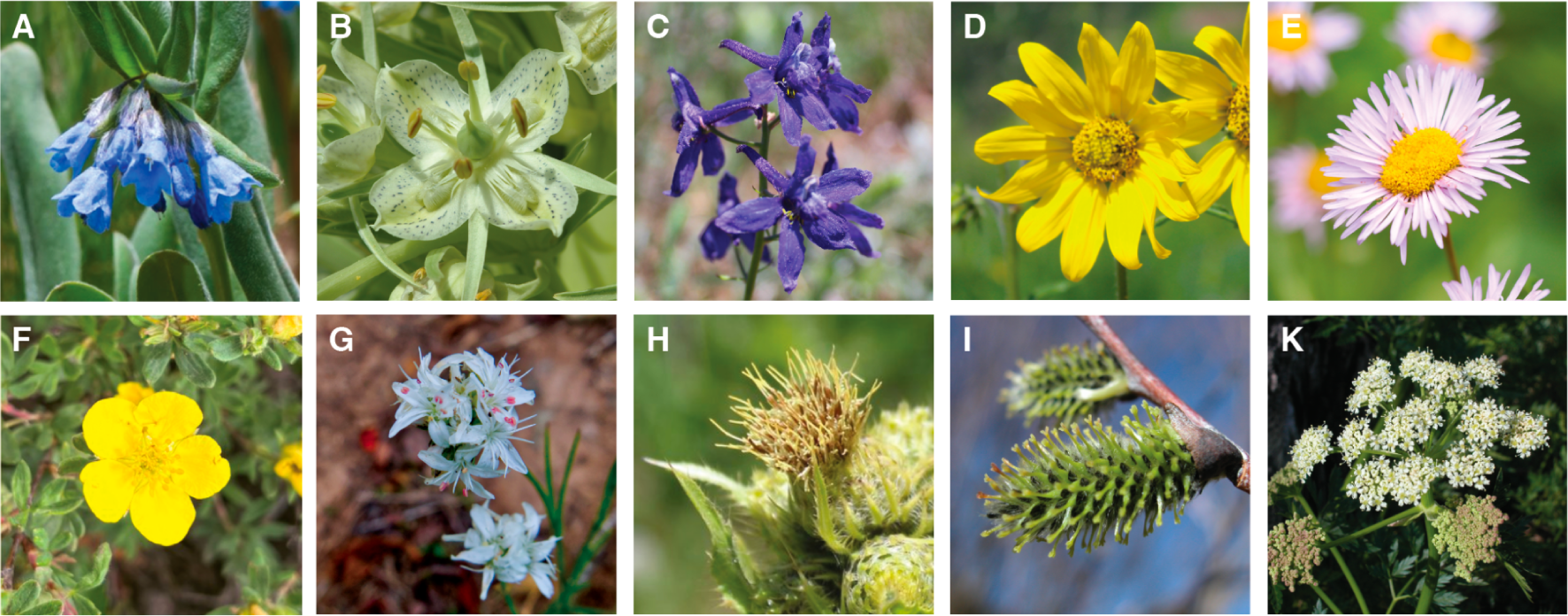
Photos of plant species included as examples in Table 1. A. *Mertensia fusiformis*, B. *Frasera speciosa*, C. *Delphinium nuttallianum*, D. *Helianthella quinquinervis*, E. *Erigeron speciosus*, F. *Dasiphora fruticosa*, G. *Eremogone congesta*, H. *Cirsium parryi*, I. *Salix wolfii*, K. *Heracleum maximum*. Photo credits: Paul CaraDonna (A, C–F, H), David Inouye (B, I, K), and Gwen Kirschke (G).

### 1.2 Overview of Nectar Sampling and Measurement Techniques

Common techniques to collect nectar include the direct use of microcapillary tubes, filter paper wicks, or syringes (*reviewed* in Kearns & Inouye, 1993; Power et al., 2018). If nectar is scarce or difficult to access, distilled water washes or centrifugation techniques can also be used (Kearns & Inouye, 1993; Silva et al., 2004). Each technique offers different benefits and challenges, but the methods we describe hereafter use a combination of direct microcapillary tube collection and centrifugation to sample species with a wide range of floral morphologies. These techniques allow us to extract nectar volumes as low as 0.05 μl in the field.

**Table 1.**
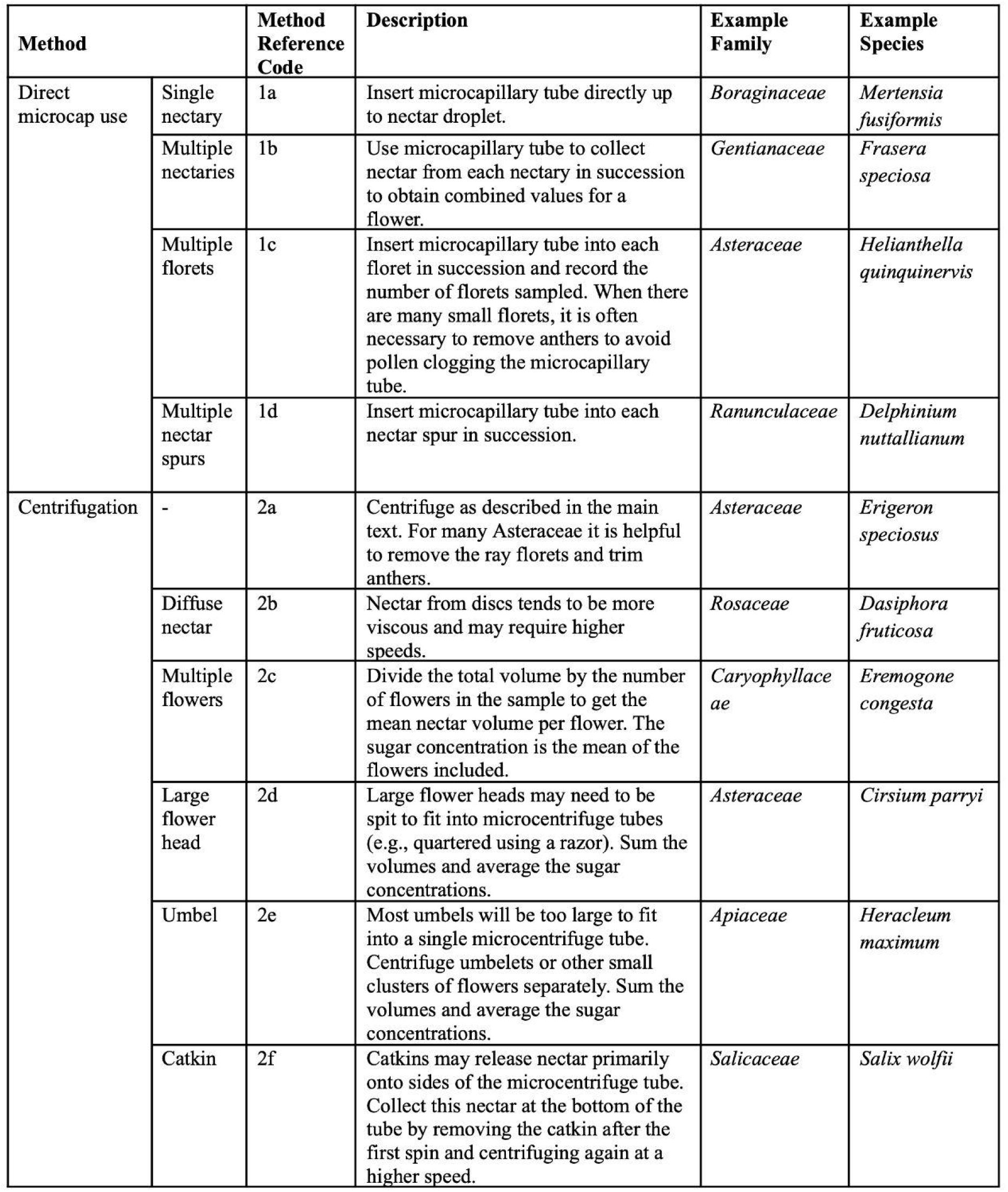
Sampling methods used to extract floral nectar and some example plant species for which they are applicable.

Our primary objective is to outline practical methods to quantify the most basic nectar traits for many species in a community under field conditions (Figure 1). It is worth noting that other nectar traits may be relevant for research questions, including sugar composition (Nicolson, 2022), amino acids (e.g., Power et al., 2018), secondary compounds (Zioga et al., 2020), pesticides (e.g., Brewer et al., 2025), and microbes (e.g., Martin, Schaeffer & Fukami, 2022). However, most of these analyses cannot be conducted in the field and often require nectar volumes larger than those obtained from a single flower.

### 1.2 Focal Study System

We sampled nectar from flowering plants in a high elevation ecosystem (2891 m asl) near the Rocky Mountain Biological Laboratory in Gothic, Colorado, USA (38°57.5′N, 106°59.3′W) over an eight year period (2018–2025). Within this montane study system, we sampled floral nectar from 53 plant species across 20 plant families with varied floral morphology, taxonomy, flowering phenology, and nectar traits (Figure 2; Table S1). The study area is a mosaic of dry and wet meadow habitats intermixed with aspen and conifer forests (Langenheim, 1962). Overall, the ecosystem is relatively water-limited and includes a wide range of flowering plants and flower structures, including many that are considered challenging to sample from (e.g., Asteraceae, Apiaceae; small and clustered floral morphologies, and flowers with small amounts of nectar), making it a useful model system to demonstrate the general utility of the nectar collection methods. While a montane ecosystem was our floral nectar sampling context, the methods described here are applicable and can be adapted to many other systems.

### 1.3 Results of Floral Nectar Variation Across 53 Species

Across the 53 species sampled, nectar volume for flowers with nectar ranged from 0.021–55 μl (median 0.69 μl) per flower ; nectar sugar concentration ranged from 0.051–1.2 mg/μl (median 0.61 mg/μl); and total sugar content per flower ranged from 0.0043–49 mg (median 0.39 mg) (Figure 1, Table S1). Among the species in this ecosystem, many flowers commonly have low nectar volumes (< 1 μl) and relatively high sugar concentrations (> 0.6 mg/μl). All 53 plant species produced floral nectar, but for 16 species, we encountered individual flowers with no measurable nectar.

## 2. Applying Nectar Collection Methods Across Space and Time: Planning and Workflow

### 2.1 Planning

Nectar sampling from many species across the season requires careful planning and organization; while considering these factors simply constitutes the effective planning and organization required for any ecological study, there are details that took enough trial and error that we considered them worth unpacking in the supplement (see A1). Useful considerations to make the nectar sampling workflow efficient and effective include using natural history knowledge, field observations, and existing data resources to organize the target plant species list by flowering time, habitat, rarity, and sampling method; choosing whether to collect the ambient quantity of nectar (standing crop) or nectar from bagged flowers; and deciding whether to control for biotic and abiotic factors influencing nectar production and evaporation.

#### Sample Size

Sample sizes of 10-30 flowers per species appeared to capture natural variation, although sample sizes of 10 or fewer are regularly included in nectar analyses (Michelot-Antalik et al., 2025; Prys-Jones & Corbet, 2011). For uncommon or difficult to sample species, even a single sample can be valuable. When sampling across a plant community, there is a tradeoff between sample sizes and the number of species one can include. One should also consider the unit of replication relevant to the research question (e.g., flowers vs. plants, sampling dates, sites).

### 2.2 Obtaining Nectar Samples in the Field

We use two general methods to extract nectar from flowers: microcapillary tubes (Method 1), and centrifugation (Method 2). Each method was used for about 50% of the species we sampled: Table 1 provides a breakdown of variations on these two methods for use with different floral structures; Figure 3 is a decision tree to help select a method, given relevant floral traits; and Appendix 2 contains extended descriptions of both methods.

**Figure 3.**
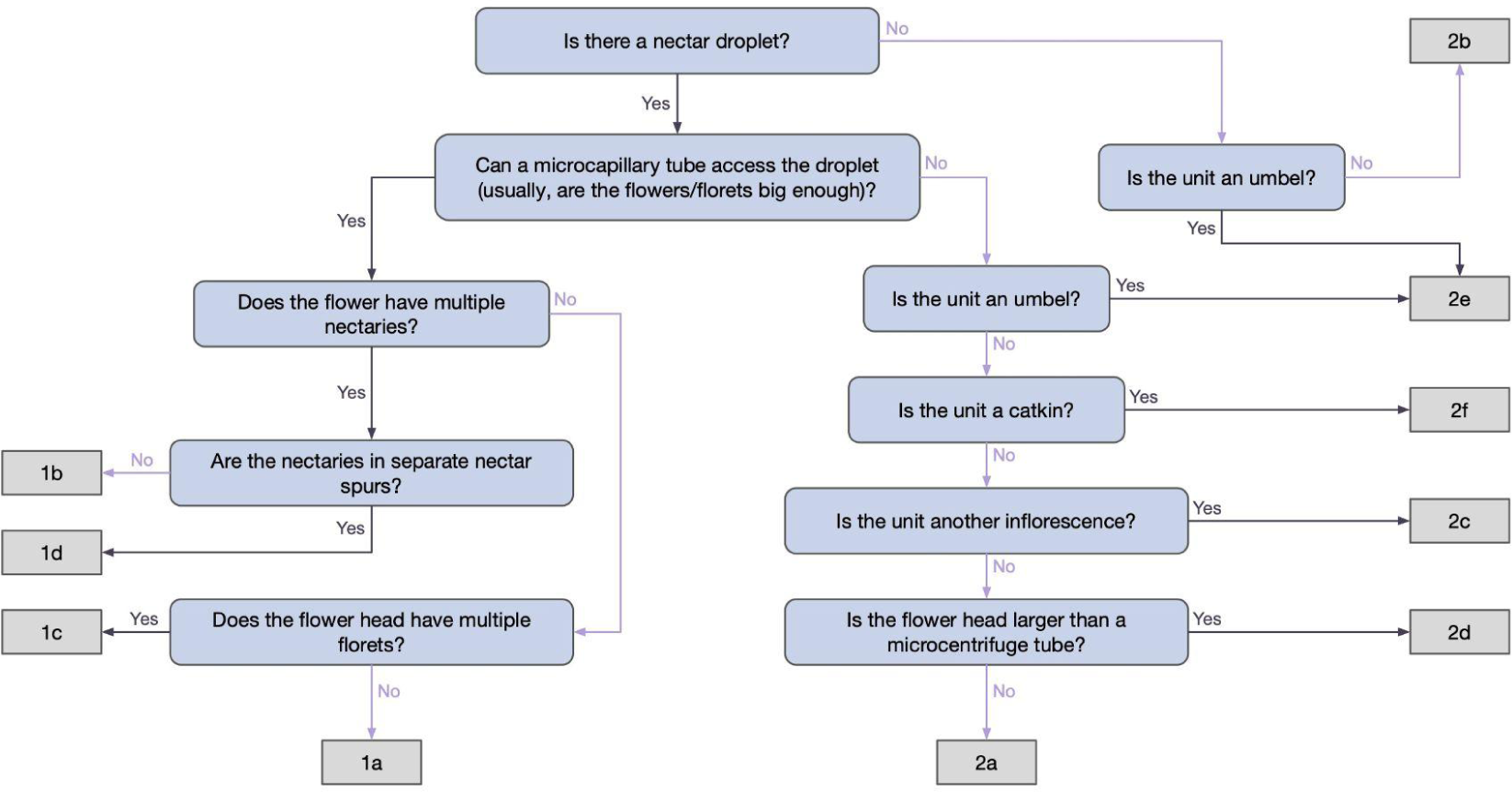
Decision tree for which nectar collection method to use for different plant species. Number-letter combinations refer to sampling methods in Table 1.

#### Method 1: Nectar collection via microcapillary tubes

Microcapillary tubes are used to extract nectar directly from the inflorescence by placing the tip of the tube against the nectar droplet(s). This should be repeated for all nectaries (Figure 4 A-C).

**Figure 4.**
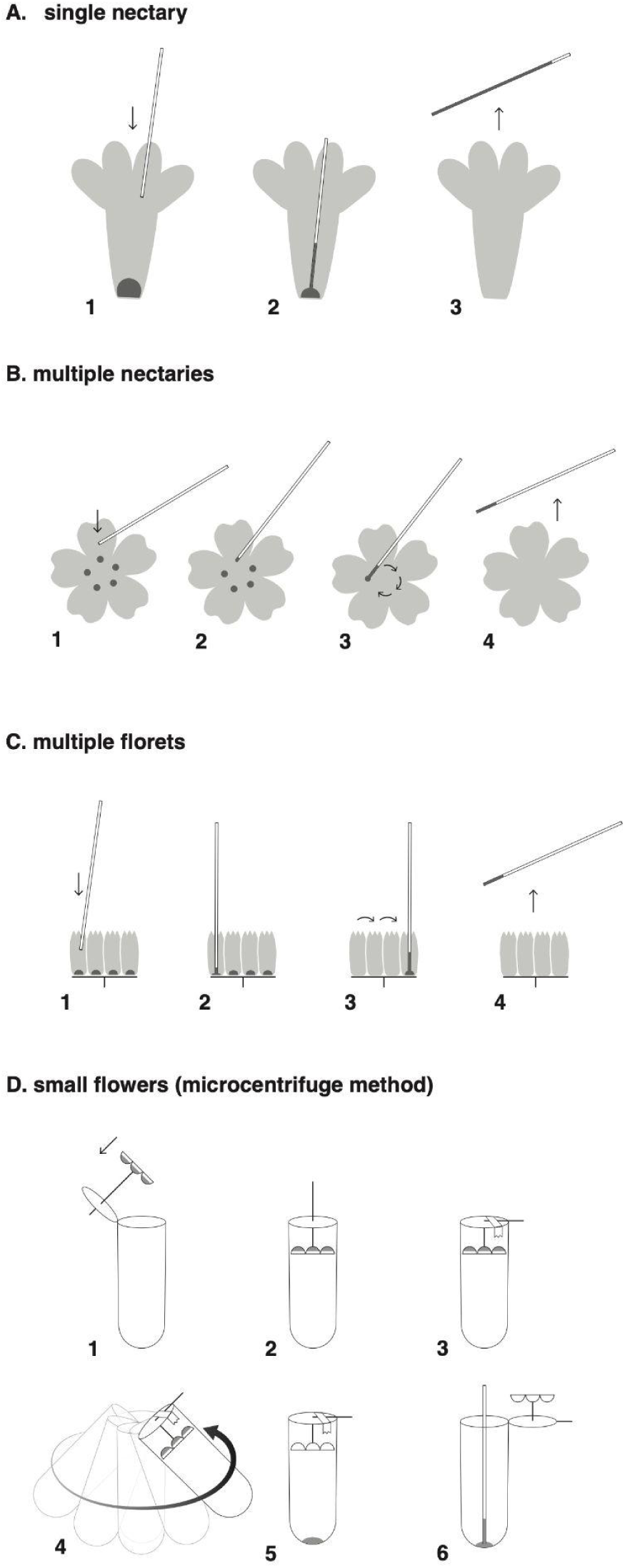
Extracting nectar. Illustrations A-C depict extraction of nectar directly from common floral structures: (A) flowers with a single nectary or nectar droplet, (B) flowers with multiple nectaries, and (C) flowers with multiple florets. The general procedure is as follows: first, the end of the microcapillary tube is placed near the nectary, touching the nectar droplet (step 1); next, the nectar is drawn into the tube via capillary action (step 2); this procedure is repeated for other nectaries or florets if needed until all nectar has been extracted; sampling is then complete and the sample is ready for volume and concentration measurements (steps 3 and 4). Illustration D depicts nectar extraction by centrifugation. (1) The stem of a flower is threaded through a hole made in the lid of a microcentrifuge tube, (2) the tube is closed with the flower head at the top, (3) the stem is secured to the outside of the lid with tape, (4) the centrifuge tubes are spun, (5) a nectar droplet collects at the bottom of the tube, and (6) the droplet is collected with a microcapillary tube. For a photographic version, see figure S1. Design and illustration by Paul CaraDonna and Richard Perenyi.

#### Method 2: Centrifugation

Flowers are transported back from the field in a cooler. Holes are punched in the lids of microcentrifuge tubes and the pedicle or peduncle of the flower is threaded through to attach the flower to the inside of the lid. The tubes are then spun until the nectar collects at the bottom of the tube, where it can then be collected with a microcapillary tube (Figures 4 D and S1).

Once the nectar is in a microcapillary tube, the volume can be assessed by measuring the portion of the tube filled, and the concentration can be measured in the field using handheld refractometers.

##### Box 1.

Nectar sampling toolbox: what to bring to the field for nectar sampling. For efficient and organized sampling in the field, we have put together nectar sampling toolboxes. Our nectar toolboxes include the following supplies which are necessary for nectar collection: (A) small-volume sugar refractometers for low and high concentration ranges; (B) garden twist ties; (C) hand lens (10x); (D) small water dropper and cleaning cloth; (E) 15 cm ruler; (F) small pollinator exclusion bags; (G) large pollinator exclusion bags; (H) pocket knife; (I) assorted microcapillary tubes (shown here: 1 μl, 2 μl, 5 μl); and (J) fine tip forceps. Other general items not included: pencils, datasheets, and field notebook. All images are to scale.

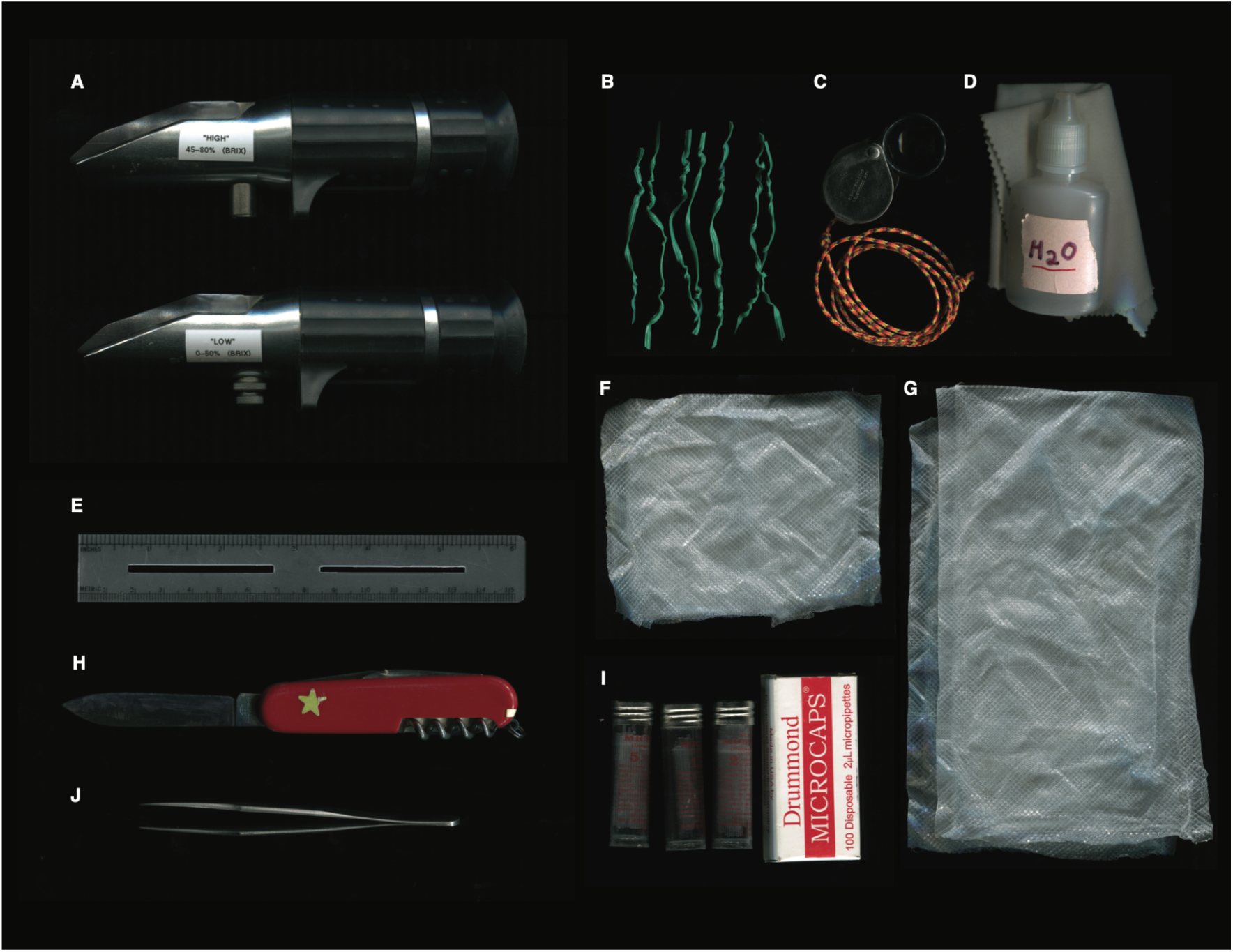

## 3. Troubleshooting Guide To Common Nectar Sampling Problems

The methods detailed above work well to extract and quantify the nectar traits of a wide diversity of flowering plants (Figure 1), but we nevertheless encountered challenges sampling nectar in the field. Therefore, we provide a troubleshooting guide which can be applied to many other species in other ecosystems.

### 3.1 Flowers with nectar spurs or long-tubed corollas are too narrow for microcapillary tubes

Flowers with nectar spurs or long-tubed corollas are often not ideal candidates for centrifugation, since they may be fragile and/or have morphologies that obstruct the exit of the nectar from the flower during centrifugation. Assuming destructive flower sampling is acceptable for your project, the tip of the microcapillary tube can be used to split the nectar spurs open lengthways to access the nectar. If you are attempting to keep the flowers intact, gently squeezing the tip of the nectar spur will push most of the nectar out to the wider portion of the spur, where you can access it (similar to Cruden & Hermann, 1983).

### 3.2 Microcapillary tubes are clogged

A clogged tube is indicated when ample nectar is visible, but is not being drawn into the microcapillary tube. Common culprits are: (i) particulates, such as pollen and dust, and (ii) high viscosity nectar.

#### Microcapillary tubes are clogged with particulates

Clogging with pollen grains can be preempted by removing anthers from flowers before sampling. If a microcapillary tube is clogged and the sample is valuable, it can be carefully broken on one side of the blockage to allow a portion of the nectar to be released onto the refractometer to obtain a sugar concentration reading without a volume measurement (make sure not to drop any small fragments of glass, and to avoid scraping oneself or the refractometer with the broken end). When centrifuging, filters that fit inside the centrifuge tube may also be used, although this increases expense (Knabe et al., 2015).

#### Microcapillary tubes are clogged with high viscosity nectar

Using a microcapillary tube with a larger diameter (usually one of a higher volume with the same length) can reduce clogging with viscous nectar. For example, for *Lathyrus leucanthus* (Fabaceae), which has a mean nectar volume of ∼ 1 μl, we sometimes used a 5 μl sized microcapillary tube because the nectar was so viscous. Using a pipet bulb will increase suction (Kearns & Inouye, 1993). High viscosity nectar can also be mixed with a known volume of distilled water to make it less viscous, so long as the quantity of water added is noted to be included in calculations later (*sensu lato* Kearns & Inouye, 1993).

### 3.3 Nectar volume is greater than microcapillary tube capacity

Flowers with relatively large and variable nectar volumes can exceed the capacity of the microcapillary tube. In these cases, multiple microcapillary tubes can be used, and separate refractometer readings taken for the nectar in each microcapillary tube. We recommend not combining the samples directly on the refractometer because the first one can dry out while the second is being collected. These volumes (i.e., lengths) can be summed across tubes and an average can be taken of the nectar refractometer readings. For example, *Frasera speciosa* (Gentianaceae) produces upwards of 30 μl of nectar per flower which requires multiple microcapillary tubes to measure the entire volume of nectar in a single flower. Therefore, to obtain traits per flower, we summed the total volume of nectar across all microcapillary tubes, and took the mean refractometer reading across all samples of the same flower.

### 3.4 Nectar sample is diluted with water droplets

Nectar samples may get diluted with water from droplets on a pollinator exclusion bag or flower petals, or from a microcentrifuge tube that is not completely dry. This can be avoided by carefully checking the flowers and microcentrifuge tubes. Because the volumes of nectar measured are so small, instances of dilution can often be easily identified because they result in incredibly low sugar concentrations compared to other nectar samples (e.g. 1–4 Brix compared to 15–80 Brix).

### 3.5 Nectar samples with uncertain sugar concentrations

When working with separate refractometers for low and high concentrations, it is not always clear which one to use for a given sample. Previous measurements from the same or related plant species can be a guide (see Table S1 for our measurements); otherwise, it is necessary to proceed with some trial and error. It can also be difficult to measure nectar in species for which the sugar concentration ranges between 40-60 Brix, as they cannot consistently be read on either refractometer. If the sample is large enough, some of it can be kept in the microcapillary tube to be used on the second refractometer if needed. Such samples can also be diluted with a known volume of distilled water, as discussed above, to consistently be read on the low concentration refractometer.

### 3.6 Refractometer readings are unclear with low nectar volumes

If a single flower does not yield enough nectar to obtain a clear refractometer reading, then multiple samples can be combined within a single microcapillary tube to increase the volume of the sample. For flowers sampled directly, the additional length of nectar in the microcapillary tube after it is inserted into each flower should be noted to determine the volumes from individual flowers, even if only one concentration value can be generated from the combined sample. It is also important to check between each flower that the apparent low volume is not because the microcapillary tube is clogged (see above).

Multiple flowers can also be centrifuged together in a single microcentrifuge tube (see Table 1, method 2c). In both cases, the sugar concentration is necessarily the mean of the flowers included. Distilled water washes of a known volume may also be used in these cases; these washes capture more of the available sugars for flowers with low nectar volumes than sampling with microcapillary tubes alone (Kearns & Inouye, 1993; Morrant et al. 2009).

### 3.7 Refractometer readings are unclear in low light conditions

In low light refractometers require higher nectar volumes to obtain readings compared to sunny conditions. Overhead electrical lights or flashlights can supplement sunlight in some cases, but it is still more difficult to get readings. If possible, sample during bright sunny days and avoid sampling in the evening or on days with heavy cloud cover.

### 3.8 Refractometer readings are unclear in seemingly suitable conditions

Even under ideal conditions, it is still possible to not see clear sugar concentration readings for nectar samples. In our experience it is still unclear why this issue arises. For example, this was the case for *Mertensia fusiformis* when we had moderate volumes (e.g., 0.35 μl) and bright sunlight (GEK *pers. obs*. and Nickolas Waser, *pers. comm*.). At the moment we do not have a solution for these relatively uncommon situations.

### 3.9 Very small budget for sampling materials

While using a new microcapillary tube for each sample is considered best practice, microcapillary tubes can be cleaned for reuse to reduce cost and waste by soaking, or filling and emptying them with water or ethanol (Corbet, 2003; Curden & Hermann, 1983). The microcentrifuge tubes can be soaked in water, rinsed, and thoroughly dried for reuse. Pay particular attention to the cap of the microcentrifuge tubes when making sure no water droplets remain.

## Acknowledgements

Initial method testing was conducted by Atticus Murphy in 2018. Thank you as well to the following people who assisted with nectar collection: Mikhaela Ferguson, Elsa Godtfredsen, Elsa Youngsteadt, Gillian Churchland, Isaac Steinmann, Emilie Gitter, Nick Waser, Alissa Doucet, Natalie Fischer, Julie Donohue, and Zoe CaraDonna. Thank you to Nick Waser for guidance on the initial nectar sampling process, and to Nick Waser, Mary Price, and Amy Iler for feedback on and discussion of developing methods. The Iler-CaraDonna lab group, Elsa Youngsteadt, and Laura Taylor provided valuable feedback on this manuscript.

Funding was provided by an Agnes Scott College Advantage Award, a RMBL undergraduate scholarship, The Frances Marx Shillinglaw Women in Science Fund, Anonymous Alumna ‘64 Research Fund, the Doerpinghaus Fund, the Scion Natural Science Association, the RMBL Krakauer Endowment Fund, and a RMBL Graduate Fellowship Langenheim Award (GEK); NSF DEB 1754518, and Chicago Botanic Garden (PJC); NSF GRFP under Grant No. DGE-1842165, RMBL (JAB); RMBL Scientist Fellowship (JEO & PJC)

We thank the Rocky Mountain Biological Laboratory for providing access to facilities.

## Notes

https://doi.org/10.17605/OSF.IO/XMU3F

### Competing Interest Statement

The authors have declared no competing interest.

### Summary of Updates

Fixed error in sugar concentration panel of Figure 1; updated supplemental Table S1.

