## Supplemental Figure 1 for "A Practical Roadmap For Sampling Floral Nectar From Communities of Many Plant Species"

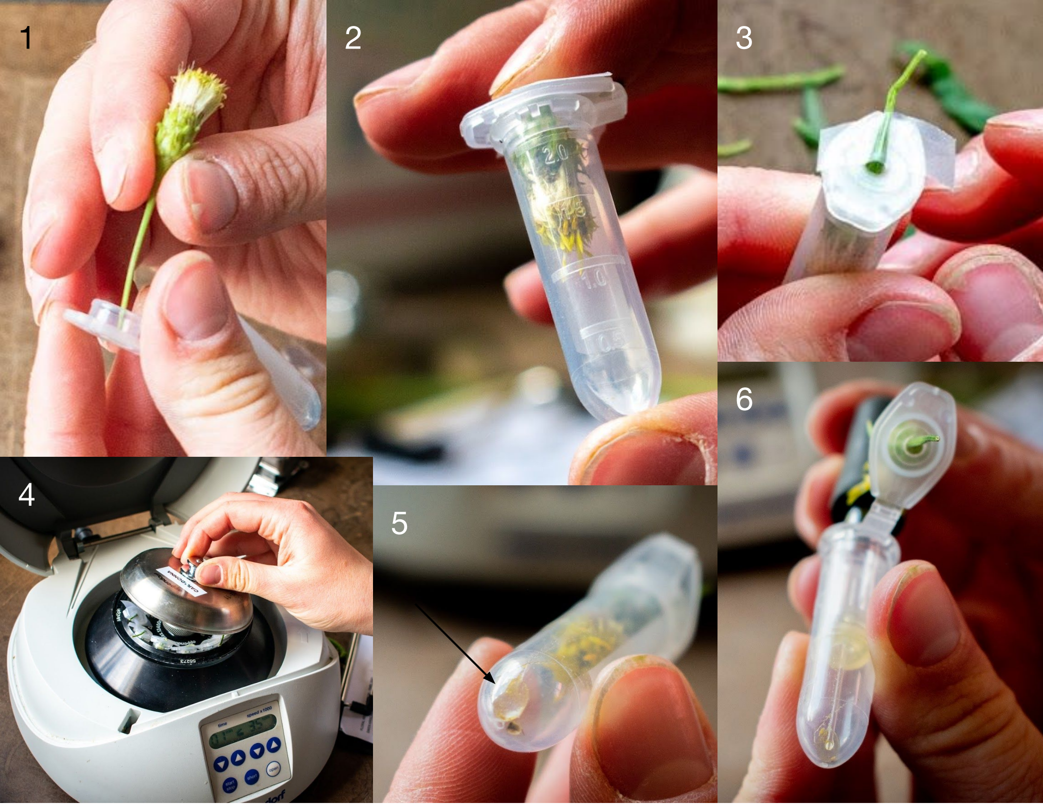


**Figure S1.** Nectar collection via centrifuge. (1) The stem of a flower is threaded through a hole made in the lid of a microcentrifuge tube, (2) the tube is closed with the flower head at the top, (3) the stem is secured to the outside of the lid with tape, (4) the centrifuge tubes are spun in a mini centrifuge, (5) a nectar droplet collects at the bottom of the tube, and (6) the droplet is collected with a microcapillary tube. All photos by Victoria DeLira.
