## Appendices for "A Practical Roadmap For Sampling Floral Nectar From Communities of Many Plant Species"

**A1: Extended Discussion of Applying Nectar Collection Methods Across Space and Time: Planning and Workflow**

### A1.1 Project Planning and Background Information Collection

The first step in nectar sampling is to determine the specific research questions and objectives which nectar sampling will address. Different questions will naturally determine which species and how many should be sampled, as well as which nectar traits should be measured. In particular, it is important to consider whether it is necessary to measure the standing crop of nectar (ambient quantity of nectar), which may be reflective of what pollinators encounter, or the nectar production capacity of plants (see Sponsler, Iverson, & Steffan-Dewenter, 2023).

Once the target plant list has been determined, it is important to review relevant natural history, morphological, and taxonomic information to help determine which sampling methods (Table 1; Figs. 1–2) are most appropriate and when and where to sample species. Particularly useful information to consider includes: flowering phenology (both dates and durations), floral morphology, local plant and flower abundance, and local spatial distribution (if not using existing sites). In the absence of local occurrence and flowering phenology data, other useful resources include: herbarium records, local and regional guidebooks, community science records (e.g., iNaturalist and Budburst), and aerial imagery. Using this information, the target plant species list can then be organized by flowering time and habitat, making note of which species have short flowering durations or are found in low abundance in the landscape. Second, target species can be grouped by their floral morphologies, which dictates the most appropriate sampling method (e.g., larger, open flowers that can be sampled easily with microcapillary tubes versus smaller clustered flowers which require centrifugation; see Section 2.2, Figure 3, and Table 1). We also suggest dissecting flowers to see where nectaries are located and nectar accumulates before selecting a nectar collection method. In some species nectaries can be in non-obvious locations (e.g., on the petals for *Hydrophyllum fendleri*). We have found that grouping species in this way greatly increases sampling efficiency and data quality.

### A1.2 Bagging to Exclude Pollinators Prior to Nectar Sampling

#### Deciding whether to bag flowers before sampling

Sampling nectar from unbagged flowers is useful to determine standing crop available to pollinators, while using pollinator exclusion bags is an attempt to standardize a replenishment period for the flowers sampled. However, nectar replenishment is stimulated by nectar removal for some plant species (e.g. Luo, Ogilvie, & Thomson, 2014; Carisio et al., 2022). Thus, measurements after a period of bagging may not represent the maximum production for these flowers over the bagging period.

#### Bag materials and methods

We place exclusion bags over flowers on plants ~ 24 hours prior to sampling to allow nectar to accumulate and to prevent pollinators from depleting it. We bag individual flowers, clusters of 2-3 flowers, or entire inflorescences, depending on the floral and inflorescence morphology of a species. We use mesh pollinator exclusion bags of various sizes, secured around stems using green garden twist ties. We have found that Midco Pollination Seed Hybridization Bags (366 micro pore diameter) work well under a variety of field conditions (UV radiation, wind, rain, snow), but other options are available (e.g., Kearns & Inouye, 1993). These bags can be used for many seasons as long as they are checked for holes and repaired as needed. For small plants that grow low to the ground, we have used modified insect nets with sand-weighted bases to cover entire plants (with 1 mm mesh; following Thomson, Forrest, & Ogilvie, 2011) because bagging individual or clusters of flowers can be challenging, time consuming, and destructive. This method is faster to deploy and covers many flowers at once. Note that when sampling from plants where many flowers were bagged together, it is important to watch or re-bag the flowers not yet sampled to prevent pollinators from visiting during sampling. To minimize stress on plant structures and keep fragile flowers intact, we propped up bags of both types using bamboo skewers and twist ties as necessary.

### A1.3 Day-to-Day Decision Making

When deciding what species to bag or sample on a given day, it is helpful to consider the following: (i) species blooming currently, based on floral counts, observations, etc.; (ii) sampling locations—we tried to limit the number of, and distance between, sites visited in one day to maximize sampling time; (iii) sampling methods—we aimed to take samples to the lab to be centrifuged in batches; and (iv) likelihood of rain—cloud cover makes refractometer readings difficult, and rain droplets can collect in some floral structures, diluting the nectar.

To determine how many flowers to bag on given day, it is useful to consider the following: (i) the overall target sample size, (ii) if all samples will be collected on one day at one site, or if they will be stratified across multiple sampling dates and locations, and (iii) other potential sources of error that may comprise sampling sizes. For example, some species consistently produce similar quantities of nectar in every flower, while in others some individual flowers do not produce any nectar. Thus, one will need to bag significantly more flowers from the latter to obtain the same number of nectar samples as from the former. This can require some trial and error, which should be included in a plan for nectar sampling.

### A1.4 Plan for Damage to Flowers from Nectar Sampling

While nectar sampling likely has little effect on plant and pollinator populations and communities, it may affect other observations and measurements if done on the same focal population. If possible, it is usually preferable to collect nectar from plants outside of study populations for other data collection, such as measurements of pollinator visitation or seed set, or to at least account for days and flowers sampled. There were some species we did not sample because there were very few at a site and we did not think we could sample them without damaging them. Non-destructive sampling, particularly in cases where multiple nectar samples are needed from the same flower, is most feasible for species with large volumes of easily accessible nectar, particularly those with sturdy floral structures (e.g. Wyatt et al., 1992).

### A1.5 Potential Factors Driving Nectar Trait Variation

Several factors are known to affect nectar production and evaporation, and depending on research objectives and study design it may be important to consider and control for them. It is also important to keep these factors in mind when applying nectar measurements from literature reviews to other systems with differing conditions. These factors include time of day, cloud cover, water availability on various time scales, pollinator visitation, and floral age and stage (e.g., from anther dehiscence to stigma receptivity in a protandrous flower) (e.g., Pleasants, 1983; Kirschke et al., in prep; Gallagher & Campbell, 2017; Waser & Price, 2016; Carroll et al., 2001; Petanidou et al., 1999; Veits et al., 2019; Luo, Ogilvie, & Thomson, 2014; Chabert et al., 2018). Finally, nectar robbing can change the rates of nectar evaporation in some floral structures, as well as result in different microbial communities (Pleasants, 1983; Martin, Schaeffer & Fukami, 2022). While we advocate for keeping these factors in mind, we have found nectar data to be informative despite such limitations, and thus do not think that nectar collection should be avoided entirely due to this variation.

**A2: Extended Descriptions of Nectar Sampling Methods**

***A2.1 Nectar Extraction***

*Method 1: Nectar collection via microcapillary tubes.* For larger flowers, those with nectar spurs, and/or visible nectar droplets, a microcapillary tube can be used to extract nectar. This method works well for a wide range of flowers, and nectar volumes from < 1 μl to >40 μl). Nectar collection via microcapillary tubes is one of the most common and versatile methods of nectar extraction (e.g., Baude et al. 2016; Gallagher & Campbell, 2016; Zhang et al., 2023). However, this method may underestimate sugar concentration in some species because sugars may be concentrated near the nectaries and not completely collected by capillary action (Petit et al., 2011). To extract nectar using this method, a microcapillary tube (e.g. Drummond MicroCapsⓇ; must be evenly bored, not tapered) is inserted into a flower and placed against the nectar droplet. The nectar droplet is drawn into the microcapillary tube via capillary action (Figure 4 A-C; Kearns & Inouye, 1993). The nectar in the microcapillary tube is easily seen as a clear liquid. With the nectar in the microcapillary tube, volume and sugar concentration can be measured (see *A2.2* *Determining Nectar Volume and Sugar Concentration*).

Although sampling with microcapillary tubes is usually straightforward, it still requires adjustments for each species, such as sampling from multiple nectaries per flower or removing anthers to prevent pollen from clogging the microcapillary tubes; for a guide to common variations, see Table 1 and *3.* *Troubleshooting Guide To Common Nectar Sampling Problems*. While some groups have had success interpolating nectar traits from floral morphology and taxonomy (see Tew et al., 2022; Baude et al., 2016), we did not detect strong morphological or taxonomic signals in our data (GEK and PJC, *unpublished analysis*). Thus, sampling new species usually requires some trial and error. When deciding which size microcapillary tube to use, start with one that can contain a small volume and adjust from there, aiming for the smallest volume of microcapillary tube reasonable for the nectar volume of a given species to improve accuracy. For example, if volumes are between one and two microliters, use a two microliter microcapillary tube, rather than five microliter tube. Starting with a small microcapillary tube ensures that you can accurately measure a small volume (if you get one).

*Method 2: Centrifugation.* Although nectar collection via microcapillary tubes is effective for many species, it does not work well for species with small flowers or florets (e.g., many Asteraceae), those with nectar discs (e.g., Apiaceae), and those with non-obvious nectaries (e.g., *Dasiphora fruticosa*). For species with these floral traits, nectar can instead be extracted using centrifugation (*sensu* Kearns & Inouye, 1993; Silva et al., 2004). Nectar collection by centrifugation has three steps: flower acquisition, centrifugation, and sample extraction. We first collect flowers in the field, store them in a cooler out of direct sunlight, and transport them back to the centrifuge within 1–2 hours. If a lab facility, or any indoor space with electricity (e.g., motel room, cabin, etc.), are not located nearby, a mini centrifuge can also be run using a power bank in the field or in the back of a vehicle. Once at the centrifuge, we place the flowers in customized microcentrifuge tubes, where we have punched a hole in the center of the lid from the inside using a nail, and trimmed off excess plastic from the hole with a razor blade. The pedicle or peduncle of the flower is threaded through this hole from the inside, situating the flower (or inflorescence) on the inside of the lid. The microcentrifuge tube is then closed, and the stem is folded over the outside of the lid and secured with clear tape. These tubes are centrifuged at g-forces between 600–2400 x g for 1 minute. We use an Eppendorf MiniSpin (Catalog No. 022620100l); on this machine such g-forces require a rotation speed of 3000-6000 rpm. The rotation speed to achieve similar g-forces will vary based on the radius of the centrifuge. The g-force and duration can differ between flowers with different structures and nectar viscosities. Higher g-forces tend to cause more fragile flowers to disintegrate, but are more effective for viscous nectar such as that on nectar discs. We have found that a g-force of 810 x g (3500 rpm on the MiniSpin) works well for most species in our study system, with lower forces for more fragile structures (e.g., *Salix*), and 1300 x g (4500 rpm on the MiniSpin) for species with nectar discs (e.g., *Dasiphora fruticosa*, table S1). In instances where nectar is distributed on the sides of tubes, such as occurs with willow catkins, 2400 x g (6000 rpm on the MiniSpin) is a suitable force to accumulate a nectar droplet at the bottom of the tube after removing the catkin. Once there is a droplet, it can be collected from the bottom of the centrifuge tube using a microcapillary tube (see Figures 4 D and S1), and the volume and concentration measured as described below.

### A2.2 Determining Nectar Volume and Sugar Concentration

*Quantifying nectar volume.* Once nectar is obtained by either of the two methods listed above, nectar volume can be determined using the following calculation (Kearns & Inouye, 1993).

$$nectar volume = \frac{length of microcapillary tube filled with nectar}{total length of microcapillary tube} ✕ max volume of microcapillary tube$$

For example, if the nectar from a flower filled 24 mm of a 32 mm long, 2 μl capacity microcapillary tube, it would have a volume of 1.5 μl.

*Quantifying nectar sugar concentration.* Sugar concentration can be measured by releasing the nectar sample onto a low volume optical refractometer. We use a Bellingham + Stanley, Eclipse (product codes 45-81 / 45-82), for sugar concentration ranges between 0-50 Brix (0-0.615 mg/μl) and 45-80 Brix (0.541-1.129 mg/μl) respectively. To our knowledge this is the only refractometer currently sold that is effective for nectar samples at low volumes (< 20 μl). With practice, we have been capable of consistently reading samples of 0.05 μl, with some refractometer readings from as little as 0.025 μl of nectar. Automatic temperature calibration is not available on these models; this was not a major concern for our observational study, as the variation introduced by temperature was well below the inter- and intra-specific variation we observed. However, for experimental projects it may be necessary to factor in time for manual calibration. Although this method can have limitations when compared to laboratory methods, the lower time and expense increases the ability to sample more species, which is valuable given how little data exists on nectar traits.

Refractometer readings in degrees Brix can be converted to mg/μl as follows, where *b* is the concentration in ˚Brix (Prys-Jones & Corbet, 2011):

$$c=\frac{(0.0037921b + 0.0000178b^{2} + 0.9988603) ✕ b}{100}$$

For example, if a nectar sample has a refractometer reading of 65 ˚Brix, it has a sugar concentration of 0.858 mg/μl.

The mass of sugar contained in the sample can be simply calculated by multiplying the volume by the concentration. If the last two examples were from the same flower, with a volume of 1.5 μl and a sugar concentration of 0.858 mg/μl, this sample would contain 1.288 mg of sugar.
