## Supplemental Table 1 for "A Practical Roadmap For Sampling Floral Nectar From Communities of Many Plant Species": Table_S1_legend.rtf

Table S1. Mean, median, minimum, and maximum values of floral nectar volume, sugar concentration, and total sugar for 53 species sampled near the Rocky Mountain Biological Laboratory in 2018-2025. The general sampling method used and the specific method reference code, referring to Table 1, are also included. Note that in addition to the listed method of centrifugation,Taraxacum officinale was also sampled directly (method reference code 1c). However, this method was time consuming, and it was difficult to keep track of which florets had been sampled, so direct sampling is not recommended for Taraxacum.
